# Moving towards more holistic machine learning-based approaches for classification problems in animal studies

**DOI:** 10.1101/2024.10.18.618969

**Authors:** Charlotte Christensen, André C. Ferreira, Wismer Cherono, Maria Maximiadi, Brendah Nyaguthii, Mina Ogino, Daniel Herrera C, Damien R. Farine

## Abstract

Machine-learning (ML) is revolutionizing field and laboratory studies of animals. However, a challenge when deploying ML for classification tasks is ensuring the models are reliable. Currently, we evaluate models using performance metrics (e.g., precision, recall, F1), but these can overlook the ultimate aim, which is not the outputs themselves (e.g. detected species or individual identities, or behaviour) but their incorporation for hypothesis testing. As improving performance metrics has diminishing returns, particularly when data are inherently noisy (as human-labelled, animal-based data often are), researchers are faced with the conundrum of investing more time in maximising metrics versus doing the actual research. This raises the question: how much noise can we accept in ML models? Here, we start by describing an under-reported factor that can cause metrics to underestimate model performance. Specifically, ambiguity between categories or mistakes in labelling validation data produces hard ceilings that limit performance metrics. This likely widespread issue means that many models could be performing better than their metrics suggest. Next, we argue and show that imperfect models (e.g. low F1 scores) can still be useable. Using a case study on ML-identified behaviour from vulturine guineafowl accelerometer data, we first propose a simulation framework to evaluate robustness of hypothesis testing using models that make classification errors. Second, we show how to determine the utility of a model by supplementing existing performance metrics with ‘biological validations’. This involves applying ML models to unlabelled data and using the models’ outputs to test hypotheses for which we can anticipate the outcome. Together, we show that effects sizes and expected biological patterns can be detected even when performance metrics are relatively low (e.g., F1: 60-70%). In doing so, we provide a roadmap for validation approaches of ML classification models tailored to research in animal behaviour, and other fields with noisy, biological data.

**Highlights:** Evaluating machine learning (ML) models must go beyond performance metrics

Mislabels in validation data leads to underestimation of model’s performance

Underestimated metrics can cause research delays despite models being useful

We propose simulations and biological validations to evaluate model performance

Models with low standard metrics can still be powerful for hypothesis testing

## Introduction

Advances in technology have dramatically enhanced our capacity to gather data on animals (Kays *et al*. 2015; Caravaggi *et al*. 2017), giving us insight into where animals go (e.g., GPS: Cagnacci *et al*. 2010), what they are doing (e.g., tri-axial accelerometers: Brown *et al*. 2013) and with whom they interact with (e.g., proximity tags: Ryder *et al*. 2012). This drastic increase in data collection has also been accompanied with advances in computer science, namely in machine learning (ML), allowing us to process, categorise and analyse large amounts of complex data. For example, ML has been used to classify species of animals from millions of pictures from camera traps in a fraction of the time of a human performing the same task (Willi *et al*. 2019; Norouzzadeh *et al*. 2021; Whytock *et al*. 2021). Similarly ML-based classification has been used to identify individuals (Schneider, Taylor & Kremer 2022), detect behavioural states (Riaboff *et al*. 2022; Aulsebrook, Jacques-Hamilton & Kempenaers 2024) and capture social associations (Valletta *et al*. 2017; Smith & Pinter-Wollman 2021), tremendously increasing the data available to address fundamental questions in behavioural ecology. ML is particularly powerful in the context of data collection as it allows us to solve classification tasks that turn raw data into interpretable data for posterior hypothesis testing. For instance, in a study on flatback turtles (*Natator depressus*), multi-sensor data from tags was classified into distinct behaviours, including foraging. These foraging predictions were then used in a regression model to show how turtles feed more at high tide to access intertidal food resources (Hounslow *et al*. 2023). Nevertheless, for all the published studies that successfully used ML models for classification problems, there are potentially many others where low model performance (e.g. low F1 scores) made the results less appealing for publication. This has likely driven a publication bias comprising of artificially high model performance and an over-optimism about ML performance (e.g., Jennions & Moeller 2002; Saidi, Dasarathy & Berisha 2024), as well as substantial loss of research time seeking high model performance that seems unattainable.

Low scores on performance metrics (see: Rainio, Teuho & Klén 2024 for a review on commonly used evaluation metrics) understandably lead researchers to invest in a time-consuming process of ML model refinements before getting to hypothesis testing. These refinements can include trying different hyper-parameters, model architectures, or collecting additional training data. However, the improvements yielded through what is effectively trial-and-error exploration are typically marginal (Probst, Boulesteix & Bischl 2019; Sipper 2022). Increasing the training dataset is usually the most effective way of improving models’ performance, which can motivate a large time and/or financial investment. Collecting more data is, however, often not viable either due to the data collection period being limited in space and time (e.g., within a delineated field season: Shuert, Pomeroy & Twiss 2018; Christensen *et al*. 2023) or because certain biological phenomena or species are hard to observe making it challenging to collect more data (Hounslow *et al*. 2019; Zhong *et al*. 2021). Moreover, even if more data are collected, the collation and annotation of training data is time and resource intensive (e.g., labelling photos of animals with the species names: Norouzzadeh *et al*. 2021; or labelling hours video second-by-second: Christensen et al., 2023). Improving model performance can therefore be a frustrating and time-consuming endeavour.

We propose that more effective use of researcher time can be achieved by better understanding the sources of model (apparent) errors, how classification errors made by the model affect (or not) posterior hypothesis testing, and how models perform when deployed to realistic empirical tests. When comparing different methods and tweaking parameters fails to make the desired improvements, poor model performance is typically ascribed to insufficient or non-representative training data or to high class similarity (e.g., being groomed while sitting, lying or standing: Christensen *et al*. 2023). In some cases, categories that are being misclassified for one another may be merged into broader categories (Ladds *et al*. 2017) without jeopardizing the initial hypotheses (e.g., if the act of being groomed rather than postural data is of importance: Christensen *et al*. 2023). However, the true blame for (apparent) low performance may not lie solely at the feet of incomplete training data or the similarity of different classes, but rather in the assumptions about our training data. While ML models are often used to aid data collection by categorising large and highly dimensional datasets into distinct categories, data are often naturally continuous. For example, classifying behaviours is especially challenging because these often include transitional states (Reyes-Ortiz *et al*. 2016; Riaboff *et al*. 2022), such as switching from feeding to moving, making it hard to delineate exactly when in time the change occurred. Subtle differences between seemingly similar categories (e.g. an animal having different intentions when raising its head for predator vigilance versus looking for the next food patch) can also make labelling ambiguous, even for experienced observers. The inherent ambiguity and noise associated with biological systems naturally introduce inaccuracies in the labelled data used to train and (importantly) evaluate ML models.

Inaccuracies in the labelling process represent an under-appreciated challenge for our ability to evaluate classification models. While it may seem logical that inaccuracies limit models from achieving maximal performance because the models have to learn from data with errors, it appears that ML models can be resilient to some mislabels (Rolnick *et al*. 2017; see also Supplementary material). Instead, what has been almost completely overlooked is that mislabels during the training process hinder our ability to *evaluate* model performance, rather than limiting the true performance of the model. This is because a model that predicts a correct label will still get penalised (i.e. reported performance in terms of accuracy, recall, F1 score, etc.) when human-assigned labels in the validation set contain errors (see Supplementary material for a worked example). Errors in the labelling, which are common in biological data, can therefore make it impossible for models to achieve perfect performance metric scores during evaluation, even if they are in fact performing very well. Thus, accurately determining a model’s performance can and should go beyond simply relying on performance metrics.

Even when models demonstrate genuine underperformance, this should not automatically preclude their application to animal data. Contrary to computer scientists that are motivated to push the boundaries of the state of the art in ML by maximizing model performance (Birhane *et al*. 2022), biologists are primarily interested in using ML models as a tool to facilitate data processing and ultimately test biologically relevant hypotheses (Hilborn & Mangel 2013). In ecology and evolution, (statistical) model performance is typically evaluated in terms of its ability to provide robust tests of hypotheses (and estimating corresponding overall effect sizes), with much less emphasis put on the ability of models to make predictions for specific data points (Houlahan *et al*. 2017). Because of the noisy nature of biological data, our hypothesis testing tools, e.g., linear regression models (Bolker *et al*. 2009) or Bayesian inference (Ellison 2004), are already built to handle (random) errors in our data (Young 2018). Our ability to statistically account for noise (in this case errors in prediction) pulls into question whether—in the context of biological hypotheses testing—performance metric maximization (e.g. maximizing F1 scores) is a useful pursuit, or whether there are more informative ways of deciding when we should stop the training process and move towards model deployment for hypothesis testing. Here we argue that while current performance metrics are useful during model training and to compare different approaches, biologists should look beyond pure metric maximization. Instead, we should ask whether a model is able to perform well enough to confront the biological hypotheses of interest with the data that the model predicted.

In this paper, we outline new perspectives and solutions on how to assess ML model performance when classifying data for the purpose of posterior hypothesis testing. As a case study, we focus on the identification of behaviour from accelerometer (ACC) data collected on a free-ranging population of vulturine guineafowl (*Acryllium vulturinum*). Combining ACC data with ML approaches to infer behaviour is a powerful way of determining the behaviour of animals without the need to observe them directly. We first use a popular ML approach (random forest models) to classify behaviours using labels assigned from videos. To better understand sources of error and how these affect model performance metrics we compare the behavioural labels from different annotators. Annotator disagreement indicates one source of error in the evaluation process that can result in the underestimation of the performance of the model for some behaviours (see Supplementary material for a worked example).

Our models, like most classification models, still produce a number of prediction mistakes that could not be directly attributed to mislabels in the validation dataset. We therefore also investigate whether we can still use our model to test our biological hypotheses of interest. First, we develop a simulation framework to evaluate whether the classification from an imperfect model can still generate useful data for hypothesis testing. In doing so, we shift the focus from model performance metrics to the contribution that the model makes towards detecting true biological effects. We highlight that this is critical for studies in the biological sciences because prediction errors will have different impacts depending on the hypothesis that is being tested. Finally, to confirm the utility of ML models, we propose an additional ‘biological validation’ procedure that uses model predictions on unlabelled data to test simple hypotheses that we know *a priori* to be supported in our study system (see also: Brewster *et al*. 2018; Wang 2019; Rast *et al*. 2020; Giese *et al*. 2021; Lok *et al*. 2023; Aulsebrook, Jacques-Hamilton & Kempenaers 2024) for other studies that used biological validations). In this way, we can evaluate the model’s ability to predict data to test biological relevant questions before using it to test the hypothesis of interest (see: Ferreira *et al*. 2020a for a similar approach in social network analyses).

## Case study methods

### Collecting, labelling, and matching video data with ACC data

We use data from a population of vulturine guineafowl that are part of a long-term project at the Mpala Research Centre, Laikipia, Kenya. Birds are fitted with solar-powered GPS and accelerometer tags (15 g Bird Solar, e-obs Digital Telemetry, Grünwald, Germany, for more details see: Papageorgiou *et al*. 2021; He *et al*. 2022) which can be remotely re-programmed to collect accelerometer and GPS data (see Table S1 for details on ACC birds). Videos of n=14 birds (individually recognisable from unique colour bands) were collected (using Panasonic HC-V777) in the field from a stationary car. At the start of the video, the date and exact GPS-time stamp were filmed to time-match the video with the ACC data. Videos were annotated by two trained observers (co-authors: *name 2 redacted*, *name 3 redacted*) using the ELAN software (version 6.4) following an agreed upon Ethogram (Table S2). The ethogram consisted of four modes of locomotion (walking, trotting, running, flying), two modes of foraging (pecking at ground, nipping at foliage), two modes of self-directed care (preening and dustbathing), resting (seated, lying or in combination with preening or pecking), and standing alert. Note that vulturine guineafowl are almost entirely terrestrial, flying only rarely. Event behaviours (e.g., social interactions) and transitions between behaviours were categorised as ‘other’ during the video annotations. We annotated 13.4 hours of video (n=258 videos).

We conducted all data processing and analyses in R (version 4.1.1; R studio 2024.04.2). After merging accelerometer data to video data using the timestamp, we visually confirmed alignment of labels and corrected misaligned labels (example of accelerometer data and label alignment: Figure S1). Some videos (n=18) did not have distinct enough accelerometer profiles to confidently confirm or correct the alignment between ACC data and annotations, and these were removed from the training and validation data. These removals, together with periods where no bird was present in the video (1.5 hours) resulted in a dataset of 11.9 hours across the 14 birds. Annotated behaviours were merged into biologically similar categories or into a category called ‘other’ if behaviours were rare (drinking, social interactions) or not of interest (transitions) (Figure S2). This resulted in 7 behaviours for training the random forest model (Table S3 for sample sizes). After matching annotated video data to 1Hz ACC data (see below), seconds that included more than one annotation were assigned the category that took up the larger proportion of that second.

### Metric calculation and random forest model fitting

We summarised each second of 20 Hz ACC data into 1 Hz data with 14 variables: static acceleration (Xst, Yst, Zst), dynamic acceleration (dyX, dyY, dyZ), raw VeDBA, smoothed VeDBA using a running mean of 2 seconds, and the first and second power spectrum densities for all three axes using Fast-Fourier transformation also at an interval of 2 seconds (adapted from: Fehlmann *et al*. 2017; Christensen *et al*. 2023). The random forest models were built the R package ‘randomForest’ using the default setting of n=500 trees and mtry=3.

We ran a random forest model with seven behaviours (Table S3). We trained the random forest model using a leave-one-individual-out (LOIO) approach (Chakravarty *et al*. 2019). This approach avoids overestimating model’s performance, which originates from autocorrelation between the training and validation data (by having data from the same individuals in both training and validation datasets) and gives a more accurate reflection of the performance of the model when generalised to data from birds that were not included in the original training dataset (Ferdinandy *et al*. 2020). During the validation procedure, for the models in which two behaviours took place within the same second, the prediction would be labelled as ‘correct’ as long as it corresponded to one of those two behaviours. We then calculated a confusion matrix was produced (Table S4) and precision, recall and F1 scores, using the standard formulas (see Supplementary material). Finally, we trained a random forest model containing all of the labelled data to allow us to classify behaviours from new data (see ‘*Biological validation can directly test the utility of imperfect models*’).

## Inter-observer disagreement illustrates imperfections in labelled data

The reliance of human labelling—the current ‘gold standard’ (Culverhouse *et al*. 2003; Hooge *et al*. 2018; Mannix *et al*. 2021) —has largely overlooked the issue of ambiguity and errors when creating the ‘ground truthed’ classification data, at least in ecology (but see examples from computer science: Sheng, Provost & Ipeirotis 2008; Yan *et al*. 2014). Here, we quantify inter-observer disagreement (mislabels) to illustrate why labelling errors create a ‘cap’ on our model performance metrics, leading us to potentially underestimate the true performance of our models. To estimate inter-observer disagreement, we used data from 62 minutes of video encompassing all 7 behaviours that were labelled by the person who wrote the ethogram (*name 1 redacted*) and one of the two trained observers (*name 2 redacted* and *name 3 redacted*). We built a confusion matrix (Supplementary Material) and calculated F1 scores with the labels from *name 1 redacted* considered as the “ground truth”. As expected, we found substantive disagreement—14.7% of the labels—between annotators (Table 1) despite all following the same ethogram (Table S2) and being trained by the same person with whom their data were compared. We might predict that behaviours with lower inter-observer agreement (as a proxy for mislabels) are more difficult to classify due to errors in the training and validation datasets. Indeed, we find a moderate positive correlation (0.57) between F1 score and the agreement between observers (Fig 1; Table 1). However, mislabels does not completely explain some low F1 scores, particularly for behaviours with relatively high concordance (e.g. ‘Preening’ has the highest concordance but only ranks 4^th^ on the model’s F1 scores; Table 1). Such low performance can be a result of low sample size or because some behaviours that are easily distinguished on video can generate very similar streams of accelerometer data (Shuert, Pomeroy & Twiss 2018; Chambers *et al*. 2021). Further, we find that both annotator disagreement and model error are class dependent (some classes are never confused, and almost all confusions occur among a limited set of classes; Table S4 and Table S5). We expect that such errors will be almost uniformly distributed in studies that classify animal-based data (especially behaviours), and that this could result in substantial under-estimations of true model performance (see Supplementary material for a worked example) and utility for hypothesis testing.

**Figure 1:**
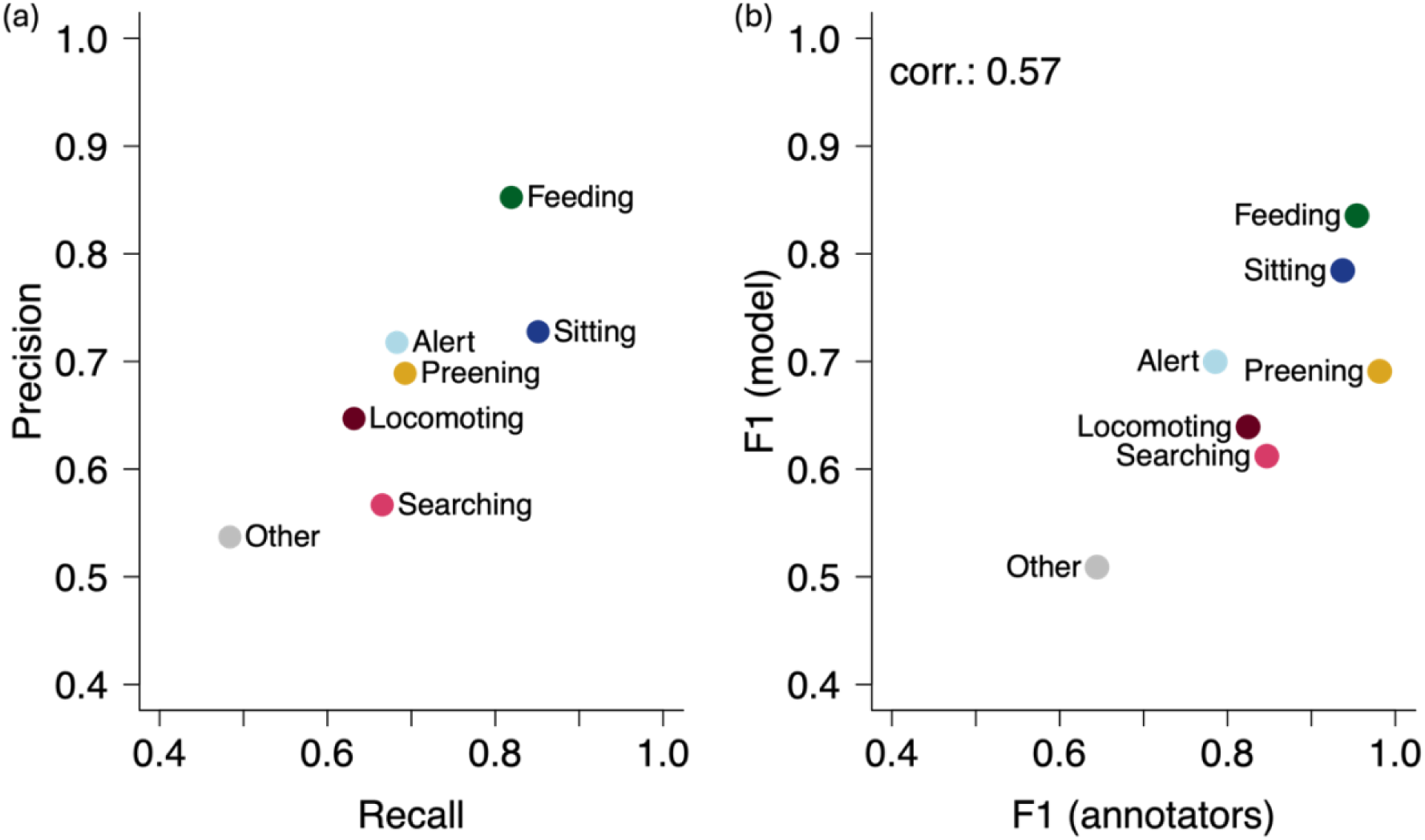
Evaluation of the performance of random forest model using standard evaluation metrics is capped by mislabels during the training process (a) Precision and recall for 7 behaviours. (b) Correlation between the F1-scores of the annotators (i.e., inter-observer agreement) and the model F1-scores (note axes start at 0.4, rather than 0.0 for easier visualisation)

**Table 1.** Labelling errors correlate moderately with the performance of ML models. Precision (model ability to minimize false positives), Recall (model ability to minimize false negatives) and F1 score (the harmonic mean of Precision and Recall) of the random forest model, and F1 score for annotators (as a measure of inter-observer agreement). Macro F1 is the average F1 score across all 7 categories.

|  | Precision | Recall | F1 (model) | F1 (annotators) |
| --- | --- | --- | --- | --- |
| Alert | 68.3 | 71.8 | 70.0 | 78.5 |
| Feeding | 81.9 | 85.2 | 83.5 | 95.4 |
| Locomoting | 63.2 | 65.0 | 63.9 | 82.4 |
| Other | 48.4 | 53.7 | 50.9 | 64.4 |
| Preening | 69.3 | 68.9 | 69.1 | 98.1 |
| Searching | 66.5 | 56.7 | 61.2 | 84.6 |
| Sitting | 85.1 | 72.8 | 78.4 | 93.7 |
| **Macro F1** |  |  | **68.1** | **85.3** |

## Imperfect models can still be powerful for hypothesis testing

Our end goal when using ML is often not the predictions themselves, but rather to use these predictions for hypothesis testing (e.g., using the categorised behaviours as a response variable in a regression). In this section, we develop a simulation framework that allows us to test the utility of ML models for hypothesis testing—specifically determining the ability to detect a given effect size, and rates of resulting false-positives and false-negatives, using a statistical model fitted to data containing imperfect predictions. We start by generating an observed second-by-second behaviour dataset that spans 10 days (collecting 7200 samples per day, e.g. 2 hours of 1 Hz accelerometer data). For each simulated second, we allocate a behaviour using the real probabilities of these behaviours taking place (based on the prevalence in the labelled guineafowl dataset), and then generate inferred behaviours by sampling from the confusion matrix (i.e. the probability that the real behaviour is inferred as each of the other behaviours) from the random forest model (Table S4). For the purpose of this simulation, we considered that the confusion matrix is not influenced by annotator disagreement in the labels (i.e. it represents “ground-truth” confusion matrix). To create a biological effect, we alter the frequency at which a specific behaviour was expressed during five of the days (e.g. increasing ‘searching’ from one time period to another by between 0 to 20% of the time budget in increments of 1.25% while reducing all other behaviours proportionately). We repeat this process 100 times for each effect size. For each study (a simulated ‘observed’ dataset), we regress the rate of the focal behaviour against treatment (period 1 vs period 2) and report the true effect (i.e. the effect size from the simulated data) and the observed effect (i.e. the effect size after classification error is introduced using the confusion matrix), as well as the P value for each model. Because misclassifications are non-randomly distributed, we expect that one possible source of false positives would be in a behaviour that is commonly mistaken with the observed behaviour (e.g. in the case of ‘searching’ the most commonly confused behaviour is ‘locomoting’; Fig. 2e). Thus, for each study we also calculated the effect on the behaviour that is most commonly mistaken with the behaviour of interest (Fig. 2; Table 2 for ‘searching’; Table S6 for remaining behaviours). We predict that misclassifications would lead the to an underestimation of the focal behaviour and that this could lead to an overestimation of the most commonly confused behaviour(s).

**Figure 2.**
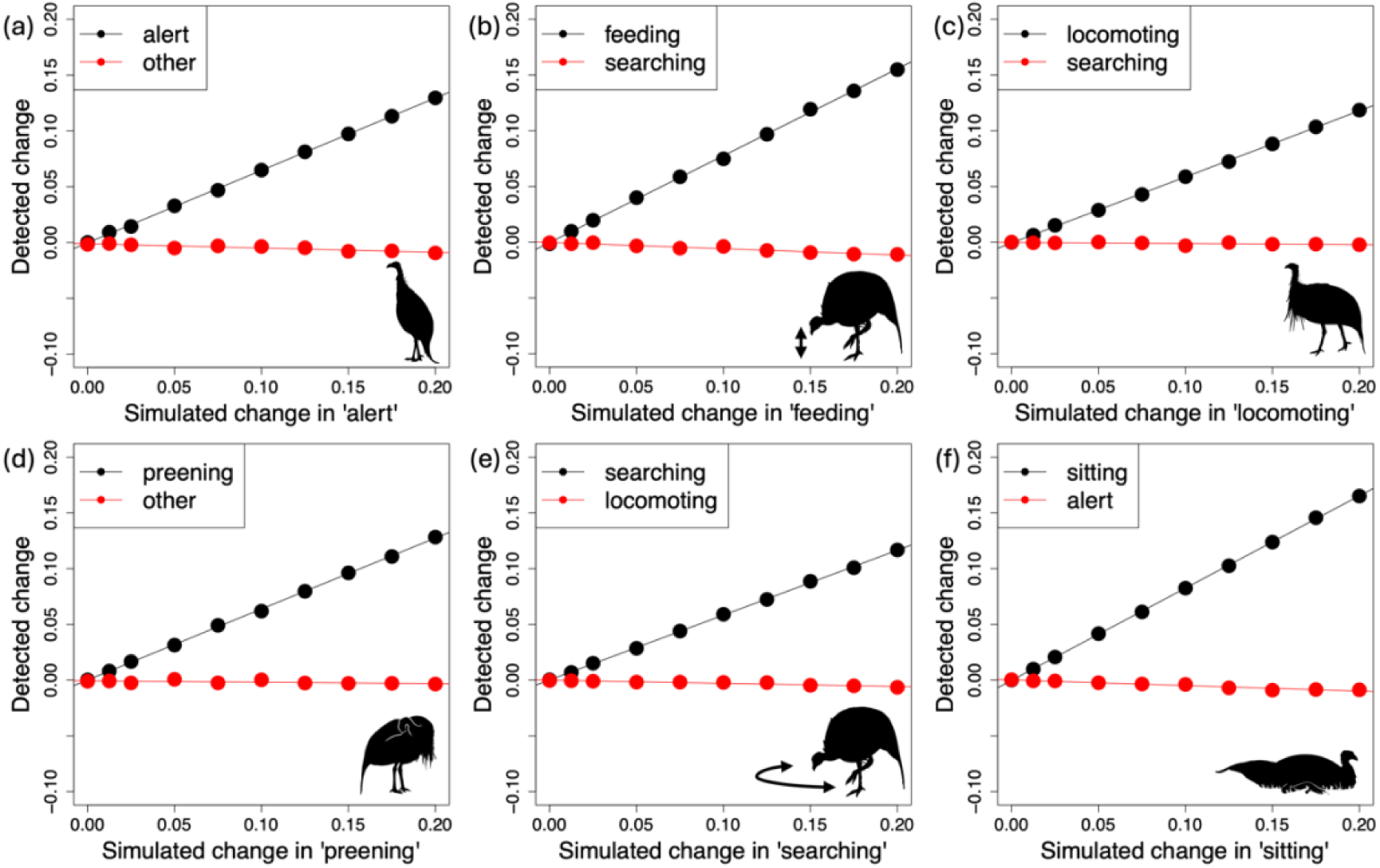
Behaviours that are increased in a dataset (‘simulated change’) can be detected despite classification error (‘detected change’). **As the simulated** frequency of a behaviour is increased (0.0 to 0.2 in increments of 0.0125; x-axis) so does the inferred rate of that behaviour from the model (y-axis, black lines and points). We find that while the frequency is underestimated, we detect no positive increase in the most frequently confused behaviour (red lines and points). Silhouettes by [name redacted].

**Table 2.** Imperfect ML models can be used to make robust biological inferences. Results from simulation where the propensity of ‘searching’ (true behaviour) is increased between two periods in the data by a given amount (simulated change, see Fig. 2e). Detected mean change captures the mean of the change between the two periods over all 100 simulated studies. Proportion of P < 0.05 is the proportion of the simulated studies that detected a statistically significant change. Detected mean change captures the mean change detected between the two periods over 100 simulations. Note that the propensity of ‘locomoting’ (confused behaviour) decreased (along with other behaviours identified in the model) to accommodate increase in ‘searching’ because the overall time available for all behaviours stays fixed, and this decrease was also detected by the model fitted to the confused data. Results for the other behaviours are presented in the Supplementary Material (Table S6).

| Simulated change | Detected mean change | Proportion of $P < 0.05$ | Detected mean change | Proportion of $P < 0.05$ |
| --- | --- | --- | --- | --- |
|  | *Searching (true behaviour)* |  | *Locomoting (confused behaviour)* |  |
| **0** | <0.001 | 0.1 | <0.001 | 0 |
| **0.0125** | 0.008 | 0.7 | <0.001 | 0 |
| **0.025** | 0.015 | 1.0 | <0.001 | 0 |
| **0.05** | 0.027 | 1.0 | -0.001 | 0 |
| **0.075** | 0.045 | 1.0 | -0.003 | 0.1 |
| **0.01** | 0.06 | 1.0 | -0.002 | 0.0 |
| **0.125** | 0.073 | 1.0 | -0.004 | 0.1 |
| **0.15** | 0.088 | 1.0 | -0.003 | 0.4 |
| **0.175** | 0.104 | 1.0 | -0.005 | 0.4 |
| **0.2** | 0.118 | 1.0 | -0.007 | 0.9 |

Our simulations show that although models produce classification errors when predicting data (in our case behaviours), the data predicted from these models can still contribute to robust biological hypothesis testing when using regressions to account for error (Table 2; Table S6; Fig. 2). We found that even small changes in behaviour can be reliably detected (i.e. high rates of significant models). Importantly, the behaviour most commonly confused with ‘searching’ (i.e., ‘locomoting’) does not increase meaningfully as ‘searching’ increases (effect sizes remain 1-2 orders of magnitude smaller than those of the real effect), suggesting that biased misclassifications (in our example) should not be expected to be a high source of false positives. Misclassified behaviours instead show a slight decrease, corresponding with the reduced time spent to make way for an increase in the focal behaviour. Further, models with no effect (simulated change = 0) also have low rates of false positives (rates of significant models is <0.1). Finally, it is worth noting that classification errors can result in underestimates of the true effect size for behaviours with low F1 scores (e.g., searching F1 = 61.2; Table 1). From our simulations, we conclude that while imperfect models can be useful, reporting the results of models should consider the relative strengths of the effects detected or report expected effect sizes beforehand that would be considered biologically meaningful (Steidl & Thomas 2001). This is a common issue in the era of big data, as very small effects can be highly significant simply because of large datasets, which is one reason for P values being unreliable (Greenland *et al*. 2016; Colquhoun 2017).

## Biological validation can directly test the utility of imperfect models

One major challenge for applications of ML in ecology and evolution is the general necessity to apply models to data collected outside of the scope of the training data. For example, we may fit accelerometer tags to new individuals during the study, use data collected after the training period has ended, or test hypotheses on data collected in different locations to those where we collected the training data. Few studies on wild animals have quantified the validity of using data collected outside of the scope of the training data (but see: Wang *et al*. 2015; Brewster *et al*. 2018; Giese *et al*. 2021; Christensen *et al*. 2023). Here we propose an approach that can both (i) test if a model allows for the identification of biologically sound patterns in true field conditions (i.e. extending the validation beyond simulation presented above), and (ii) test if a model is likely to remain useful once applied to unseen data (i.e. good generalizability). Our approach is based on biological validation, i.e., confirming that the data can identify effects known *a priori* to exist. For example, walking long distances is (in most animals) incompatible with long periods of resting or foraging. We should also expect higher rates of foraging behaviours in areas where food is abundant compared to, for example, roosting areas as animals respond to the resources in the landscape. Here we illustrate the concept of biological validation for ML models by developing four cases in which we test a “simple” hypothesis (with a known outcome) as a means of examining the performance of a model at addressing a real biological problem. While these “simple” hypotheses and *a priori* knowledge will always be species and context specific, some or all of the examples proposed here could easily be adapted to most study systems. We also note that not all behaviours may have a suitable “simple test” and that this may lead to issues of ‘coverage’ (Pei *et al*. 2017), which means not all branches of a model may be biologically validated (e.g., we have no clear prediction for the category ‘Other’ in our case study). However, we cover the behaviours of interest (i.e., that will be used to test hypotheses in future studies).

### Biological validation 1: vulturine guineafowl spend more time foraging in proximity to food supplements

To test the detection of feeding versus searching behaviours (Table S4), we take advantage of food supplementation that was available to the population between 3^rd^ June and 12^th^ July 2023 as part of another experiment in this population. Small patches of distributed millet grain (5 by 5 meters) were visited by vulturine guineafowl early in the morning. We expect that feeding behaviour should be detected more frequently in close proximity to a patch. By contrast, the easily-confounded searching behaviour (associated with slow-paced walking and scanning the ground for food) should not increase in these areas. It might even decrease, as animals have limited time (Dunbar, Korstjens & Lehmann 2009) and if they increase feeding, other behaviours should decrease. For n=8 birds, we extracted the GPS data from 06:00 to 07:30 and calculated the proximity to a foraging patch (< 15 m or > 15 m). We then use generalized linear mixed effect models (R package ‘lmerTest’) with ‘proportion behaviour per minute’ fit as a binomial response variable and proximity (< 15 or > 15 m) as a categorical fixed effect, and ‘bird ID’ and ‘Date’ as random effects. Of the total 429 hours of GPS data that satisfied the time criteria, 38 hours were within 15 m of the foraging patch with the remaining (391 hours) being outside the 15 m perimeter. The results (Table 3; Fig. 3a) confirm that guineafowl in closer proximity to the millet patch spend more time foraging. By contrast, there are very small (and negative) changes in searching behaviour.

**Figure 3.**
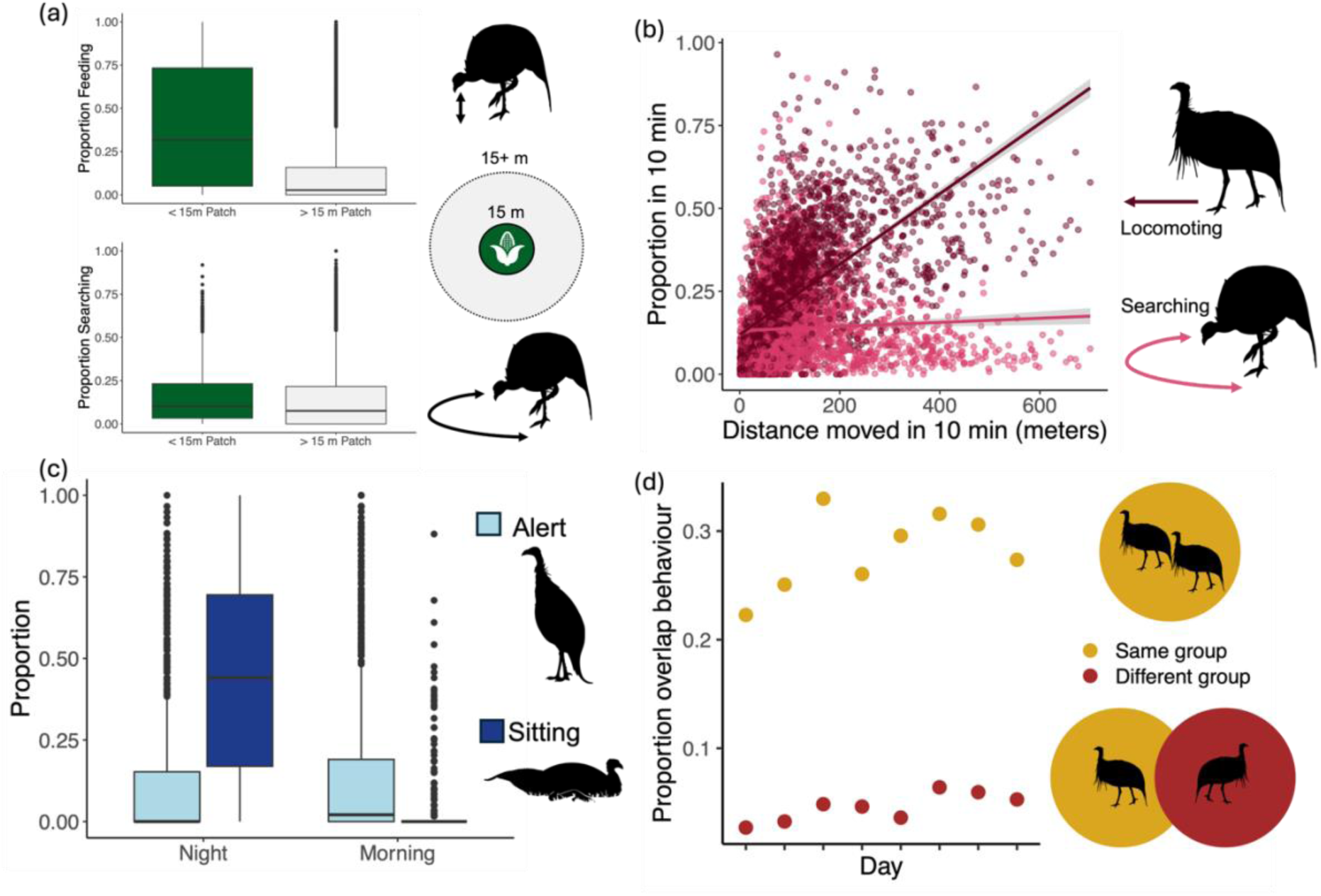
**Biological validation provides confirmation that the machine learning model is capable of testing biologically relevant hypotheses.** (a) A higher rate of feeding but not searching is detected around a supplemented millet patch. (b) A higher GPS displacement corresponds to a higher proportion locomoting but not searching. (c) A higher rate of sitting (resting behaviour) is detected at night than during the day, whereas a higher rate of alert behaviour (also immobile) is detected during the day. (d) An individual has 5 to 8 times higher overlap in second-by-second behaviour with another individual moving within the same group compared to an individual in a different group. Silhouettes by [name redacted].

**Table 3.** Biological validation demonstrating that the random forest model can detect an increase in foraging close to a known food patch. Results of a generalized linear mixed model on the inferred proportion of time spent (a) feeding and (b) searching when in proximity (<15 m) of a known food patch relative to the baseline (>15 m). Data for 8 birds across 41 unique dates with 25747 1-minute blocks.

| Predictor | Estimate | SE | z-value | *p* |
| --- | --- | --- | --- | --- |
| *(a) Feeding* |  |  |  |  |
| Patch ($<15$ m)* | 1.516 | 0.007 | 215.53 | $<0.001$ |
| *(b) Searching* |  |  |  |  |
| Patch ($<15$ m)* | -0.128 | 0.009 | -14.74 | $<0.001$ |

### Biological validation 2: vulturine guineafowl spend more time locomoting when making larger displacements

To test the ability to distinguish between locomoting and searching (Table S4), we use the GPS data to calculate the distance travelled over 10-minute periods during which the accelerometer was also collecting data. We then match this to the time spent locomoting and searching in those same 10 minutes. We expect that moving larger distances should be strongly and positively correlated with locomoting (travelling in a directed fashion). By contrast, we do not expect to find support for such a relationship for searching behaviour (travelling while looking for food). We fit generalized linear mixed effect models with proportion of the 10-minute window spent doing the behaviour of interest (‘locomoting’ or ‘searching’) as the response variable and GPS- distance travelled was the predictor variable (scaled from 0-1 using R package ‘scales’), controlling for bird ID and Date as random effects (in a binomial model). We use the whole dataset from the same periods as biological validation 1. Our analysis confirms that larger GPS displacements positively predict the proportion of ‘locomoting’ detected, but not searching (Table 4; Fig. 3b).

**Table 4.** Biological validation demonstrating that the random forest model can detect greater locomotion during periods when individuals make large displacements. Results of generalized linear mixed models on the inferred proportion of time spent (a) locomoting and (b) searching as a function of the distance travelled for 8 birds across 41 unique dates with 2830 10-minute blocks.

| Predictor | Estimate | SE | z-value | p |
| --- | --- | --- | --- | --- |
| *(a) Locomoting* |  |  |  |  |
| Distance (scaled) | 3.536 | 0.013 | 267.61 | <0.001 |
| *(b) Searching* |  |  |  |  |
| Distance (scaled) | -0.036 | 0.016 | -22.38 | <0.001 |

### Biological validation 3: vulturine guineafowl spend more time ‘sitting’ at night than during the day

To evaluate the ability to distinguish between ‘sitting’ and ‘alert’ (Table S4), we test whether night time is associated with an increase in ‘sitting’ relative to a matched day- time window. Vulturine guineafowl spend the night at communal roost sites (Papageorgiou *et al*. 2024). While some wakefulness is expected at the roost, the night should be generally associated with more ‘sitting’ (stationary) relative to morning hours when vulturine guineafowl are observed to forage with intermittent bouts of alertness. The tags of three vulturine guineafowl were reprogrammed between 25^th^ May 2023 and 6^th^ July 2023 to collect data from 18:00 to 11:00 the next day. We use these data to test whether the proportion of time spent sitting was higher between 00:00 and 02:00 (middle of the night) than 07:00 and 09:00 (morning activity hours when birds are observed to forage on the open glades), controlling for ‘bird ID’ and ‘Date’. We show that birds spend more time ‘sitting’ at night than during the morning hours (Table 5; Fig. 3c).

**Table 5.**
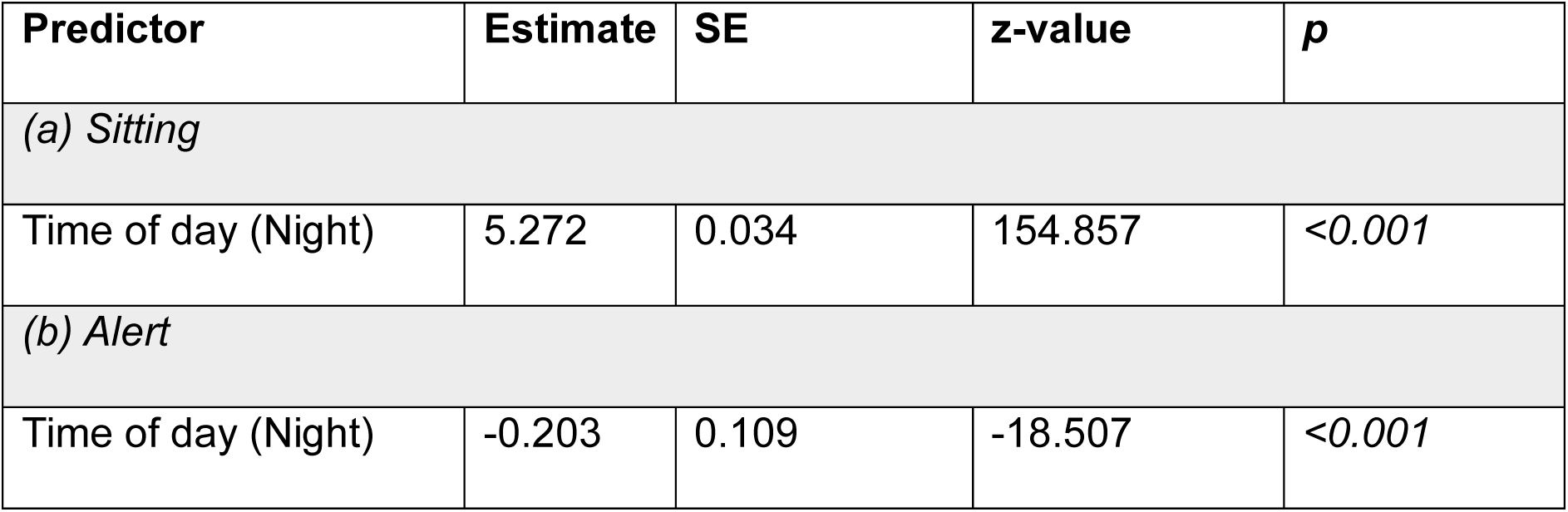
Biological validation demonstrating that the random forest model can detect changes in sitting and alertness as a function of the time of the day. Results of generalized linear mixed models on the inferred proportion of time spent (a) sitting and (b) alert at night (00:00 to 02:00) versus during the morning (07:00 to 09:00; baseline) for 3 birds across 12 unique dates with 5844 1-minute blocks.

### Biological validation 4: vulturine guineafowl show higher within-group than between-group synchrony in their activity

In some cases, *a priori* predictions about which behaviours will increase or decrease could be difficult to make, or experimental manipulations may not be feasible. In the case of group-living animals (such as the vulturine guineafowl), we can use the expectation that individuals in closer proximity (e.g. those from the same group) should be temporally more synchronised in their behaviours than dyads that are farther apart. These patterns are likely because individuals in close proximity experience more similar ecological forces, while maintaining cohesion also requires some synchronisation of activity (Davis, Crofoot & Farine 2022). with the caveat that some behaviours might also happen asynchronously, (e.g., alternating bouts of sentinel behaviour in meerkats: Santema & Clutton-Brock 2013). We use this prediction to test the performance of our model by taking the behavioural data from one vulturine guineafowl and time matching it to the data from a group member and a non-group member for 8 consecutive days (between 3^rd^ and 10^th^ of June 2022). For every second in these data, we calculated whether the behaviour was the same or not. Note that all three individuals live in close proximity, have overlapping home ranges, and experience largely identical ecological conditions. The model outputs confirm a substantially greater increase in synchronisation among individuals in the same group (mean±SD 28.2±0.04% of the time) than among individuals in different groups (0.05±0.01% of the time, Figure 3d).

## Ethical Note

The study was conducted under a research authorisation (KWS/BRP/5001) and capture permit from the Kenyan Wildlife Service (KWS/SCM/5705), the National Commission for Science, Technology and Innovation of Kenya (NACOSTI/P/21/6996), the National Environment Management Authority (NEMA/AGR/068/2017) and Wildlife Research & Training Institute (WRTI-0307-04-23). All the research was done in collaboration with the Ornithology Section of the National Museums of Kenya. Capture and GPS fitting was reviewed by the Max Planck Ethikrat Committee and the Australian National University Animal Ethics Committee (A2023/20).

Capture and handling of birds were performed by trained personnel, following methods approved by the Kenyan Wildlife Service and the Ornithology Section of the National Museums of Kenya. Birds are captured in a large, baited walk-in trap activated by a manually controlled mechanism (i.e., the trap is triggered only when it is safe to do so for the birds). After capture, they are transferred to holding cages that are covered with a tarp to minimise stress and placed in the shade. Typically, 3-4 birds per social group carry GPS tags. The total weight of backpacks (Teflon string, rubber pad) and tags combined is approximately 20.5 g, less than 2% of their body weight (females are >1280 g, males are >1440 g), which is below the generally recommended 3-5% rule (Kenward 2000). More recent evidence suggests that researchers must consider tag attachment style and animal lifestyle alongside weight (Wilson *et al*. 2021). In our case, the tags are fitted on the back, thereby fitting snugly to the centre of mass of the bird and creating limited drag during movement, unlike collars which can exert large force on the neck (Wilson *et al*. 2021). Moreover, foraging does not involve high-intensity pursuits in guineafowl (as they forage predominantly on grass, seeds and insects), meaning that the tag should not interfere with this critical activity. After capture, birds were released in groups of four to eight at a time (including tagged and untagged individuals) to avoid isolation and reduce the risk of exposure to predators. Follow-up observations confirmed that tagging had no adverse impact on birds’ ability to move, forage or stay with their social group. Table S1 contains details on the tagged animals for this study (sex and duration tags collected accelerometer data for). Tagged individuals typically carry their tag for 1-2 years, after which the tag is either removed manually during capture, the tag falls off due to the fabric wearing down or the bird is predated. Video follows of the tagged birds were conducted from a car to which the birds are habituated, keeping a minimum distance of 10 meters to avoid disturbance.

## Discussion

In this work we highlight the difficulty in accurately estimating performance metrics of ML models based on biological data, and how this can cause misguided investments or undue critiques of models. While we demonstrate this issue using behavioural data, which is ideally suited to illustrate issues of label ambiguity and its consequences for ML models, the following findings may also be more broadly applicable to other fields of ecology and evolution that use ML models to predict data for downstream analyses. We first demonstrate a moderate correlation between the ability of humans to classify data and the performance of the model at making the same classification, suggesting that errors in the validation dataset create a ceiling on metric performance. This ceiling causes an underestimation of true model performance that can have an impact on decisions of where to invest limited resources for research. Next, we suggest that while models are likely to always output a proportion of erroneous predictions, this may not necessarily hinder the utility of the model because our aim is often to test hypotheses rather than making predictions *per se*. We give an example of how simulations can be used to evaluate the utility of models for hypothesis testing, which suggest that models can still be useful even if imperfect. Finally, we propose methods for implementing validation approaches that align with the ultimate aim of using models for hypothesis testing, demonstrating that our own ‘imperfect’ models can reliably detect a number of predicted patterns in behaviour. In doing so, we provide tools for developing and evaluating models that are not only more robust but also better aligned with their intended scientific objectives.

The results of our inter-individual disagreement analysis build on the well-established finding that observers can disagree on the classification or exact temporal delineation of a behaviour (Garcia, Junior & Marino-Neto 2010; Burghardt *et al*. 2012). While rates of inter-individual disagreement do not translate directly to rates of mislabels, they show that errors in labelling are in many cases unavoidable. Some steps can be taken to improve inter-observer agreement, including extensive training, more clearly defined ethograms, shorter observation sessions to avoid fatigue and recruiting observers with long-term experience observing the animals in question (Kiddie & Collins 2014; Dai *et al*. 2020). Nevertheless, some cases are by their very nature difficult to assign, such as subtle (Dai *et al*. 2020), variably executed (Brouwers *et al*. 2023) or transitionary (Nguyen *et al*. 2021; Li *et al*. 2022) behaviours. When evaluating ML models, researchers should bear in mind the potential for mislabels, as these can penalise the reported performance metrics even if the model performs well (see Supplementary material for a worked example). Reviewing the confusion matrix and evaluating examples of incorrect predictions on the validation set can help identify any patterns in errors. For instance, errors may occur more often between transitional behaviours such as shifting from walking to running. Understanding the limitations of our human-labelled validation dataset (particularly when it comes to ‘noisy’ data) can help to set more realistic expectations for evaluation metrics and can inform decisions about when to stop training, which can avoid endlessly testing alternative models and algorithms.

Another important factor to keep in mind, is that the impact of errors in the predictions by ML-based models is not equal across all research questions. Some questions will be naturally more robust to errors, while others will be more sensitive. Imagine a ML model that is used to predict the identity of individuals interacting with one another from videos or pictures has a rate of erroneous identification (assigning the wrong identity to a detected individual). If the aim of our study was to test whether the social network of individuals is assorted by sex (Farine 2014), then even a model that makes frequent errors would still allow us to test the hypothesis robustly because assortment is very robust to imperfect sampling (Kaur *et al*. 2024) (i.e. the vast majority of edges would still accurately contribute to the test of the hypothesis). By contrast, if we wanted to instead study the spread of a contagious disease in the population, then our hypothesis may be substantially more sensitive to errors. This is because identification errors rewire the social network (Davis, Crofoot & Farine 2018). It can also artificially increase the density of edges because misclassifying individual identities can introduce connections between individuals that were never (or could never have been) observed together. The addition of just a few edges could then change the properties of spread through a social network. Thus, an error rate that could yield accurate estimations for the test of one hypothesis (network assortment) could also lead to errors for another hypothesis (e.g. dramatically overestimating the potential rate of disease transmission in the population). Similarly, some errors in behavioural predictions from accelerometery tags may be less problematic if the aim of the study is to monitor relative changes in behaviour (e.g., an increase in preening in response to higher ectoparasite load: Møller 1991). In contrast, if the exact onset of a behaviour that was previously absent is of interest (e.g., the timing of mating-related behaviours: McDonald 1989), misclassifications could make these harder to detect (or lead to false detections of such events).

In most cases, models will perform worse than hoped for on standard metric tests. One common pattern is that model performance is lower when there are more categories to classify (this is expected). In such cases, rather than testing the applicability of the ‘imperfect’ model (as presented here in our simulations), researchers are often tempted to merge categories that are frequently confused by the model and/or have similar biological functions (Ladds *et al*. 2017; Christensen *et al*. 2023) as a way of obtaining higher values in the evaluation metrics. We note that having fewer categories will almost always increase model performance across all metrics (Ladds *et al*. 2017). The reason is mathematical: a completely naïve model will have a higher probability of making a correct guess if there are fewer categories to guess from. However, merging categories does not necessarily mean that the new model is more useful. For example, combining two foraging behaviours that are often confused—such as browsing a bush and pecking at ground level—might reduce the reported error, but if these behaviours are still mistaken with other non-foraging-related behaviours that the model can predict (e.g. locomoting), then merging these behaviours might not add to any conclusion about foraging in relation to the non-foraging behaviours. Instead, merging categories for the purpose of increasing evaluation metrics may even narrow the model’s applicability and hinder hypothesis testing. For instance, merging foraging-related categories could limit our ability to test individual foraging mode/diet preferences (Toscano *et al*. 2016; Gharnit *et al*. 2022) or seasonal variation in foraging modes (Owen-Smith, Fryxell & Merrill 2010). The results from our simulations provide researchers with an additional pathway to convincingly demonstrate the applicability of their models without jeopardizing the initial research questions of interest (e.g. by losing resolution by merging categories).

High performance metrics might also be misleading and lead to overconfidence in the model. Beyond the common pitfalls that might lead to overfitting models (e.g. having the same or correlated data both in training and validation datasets; Ferreira *et al*. 2020b; Aulsebrook, Jacques-Hamilton & Kempenaers 2024), performance metrics do not always capture the different contexts in which categories might occur. For example, a binary classification model that is 99% accurate (evaluated on a balanced validation dataset) in detecting the presence vs absence of birds in a frame of a video might still produce many more false positives than actual correct predictions for the less common category (in this example, presence of birds) if the number of frames without birds is hundreds of times more common than frames with birds during the real application of the model (e.g., Silva *et al*. 2024). We argue that estimating the ability to detect effect sizes and validating models by using biological hypotheses derived using *a priori* knowledge is a more reliable method to evaluate model performance because it more closely replicates how the model will be used in the future.

Biological validation is not without its challenges. For example, we cannot give clear guidelines on how many validations to perform or guarantee that failing to detect an effect during biological validation was not because our ‘simple hypothesis’ was incorrect as opposed to the model generating too many incorrect predictions. Finally, the model might work for clear signals that rise above model classification error noise (or a delineated case, such as during an experimental treatment) but not in more complex cases, where animal responses (e.g., behavioural change) may be more subtle or masked by other simultaneous changes. For all of these reasons, validating models through simulations and testing “simple” hypotheses should not be seen as last resort to validate apparent low performing models, but as a way of obtaining greater knowledge about the future classified data, about where the strengths and weaknesses of the model may be, and about how the model (and its performance) might fit in the whole pipeline from data collection through to finalising the results (as has been proposed for statistical hypothesis testing more generally: Zuur, Ieno & Elphick 2010; Rubin & Donkin 2024).

Finally, our proposed validation approaches should not be used as a loophole to excuse poorly trained models. In the same way that reported traditional evaluation metrics can be high for low performing models (e.g. overfitted models: Bejani & Ghatee 2021), poorly trained models could output data that can appear to detect expected effect directions but totally misrepresent the effect size. The true value of validating models through simulations and hypotheses testing lies on pushing researchers to openly and critically think about the potential biases of their models and consider whether their models are vulnerable to producing misleading scientific outcomes (as discussed above), instead of evaluating them shallowly. We hope that these additional quality control layers can improve the ability for researchers to switch from model refinement to hypothesis testing while increasing the quality of ML-based proof-of-concepts works and applications. This can help mitigate the proliferation of publications featuring overfitted models, a trend already observed in other research fields (Navarro *et al*. 2021; Berisha *et al*. 2022; Saidi, Dasarathy & Berisha 2024). Biological validation adds a final layer of quality control by simulating real-world applications of the models before their publication and/or before using them to address real biological questions of interest. Such final layer of validation should elevate the standards of ML-related publication in behaviour ecology, ecology and evolution.

## Supporting information

Supplementary material (revision)

## Data availability statement

Data and code are available on Figshare: https://figshare.com/s/f015439a649d626d5aa0

## Authors contribution

CC, ACF and DRF conceptualised the study. CC and BN led the data collection of the accelerometer and video data in the field. CC wrote the ethogram and CC, WC and MM annotated the videos. DH corrected the time alignment between annotation and video data. CC trained the random forest model and performed the statistical analyses for the inter-observer agreement and the biological validations. DRF conducted the simulations. ACF collected the data and trained the deep learning models for the worked example in the Supplementary material. MO conducted the food supplementation experiment for one of the biological validations. CC and ACF wrote the first draft of the manuscript with the input of DRF. All authors read, improved and approved the final version of the manuscript.

## Acknowledgements

We would like to thank the Vulturine Guineafowl Research Programme field team: Monicah Wambui, Kennedy Kipkorir, John Wanjala and Mary Waithira Ngugi for their work in deploying tags, downloading biologging data and collecting videos that were annotated for model training. We would like to thank Mpala Research Centre for logistical support and allowing us to work on the conservancy. We would like to thank National Museums of Kenya (NMK), the National Commission for Science, Technology and Innovation (NACOSTI) and the Wildlife Research and Training Institute (WRTI) for approval of our work. CC and ACF were funded by European Research Council (grant agreement no. 850859 awarded to DRF). CC was supported by a UZH post doc grant (no. K-74312-02-01). MO received additional funding from the Swiss Federal Commission for Scholarships. DRF was funded by an Eccellenza Professorship Grant of the Swiss National Science Foundation (Grant Number PCEFP3_187058).

## Statement of inclusion

Our study brings together authors from a number of different countries, including scientists based in the country where the study was carried out.

## Data availability

Data and code will be made available on Figshare once the manuscript is accepted for publication.

