## Supplementary material (revision) for "Moving towards more holistic machine learning-based approaches for classification problems in animal studies"

**Formulas used for calculating precision, recall and F1 scores:**

Precision = True Positives / (True Positives + False Positives)

Recall = True Positives / (True Positives + False Negatives)

F1 = 2* Precision * Recall / (Precision + Recall)

**Table S1**: summary of individuals and associated accelerometer data collection period (Type denotes the dataset(s) in which the birds were included).

| Bird ID | Sex | Type | Start date | Stop date |
|---|---|---|---|---|
| W1273 | Male | Training, Millet & GPS validation | 14-02-2022 | 01-11-2023 |
| W1307 | Male | Training, Millet & GPS, Synchronisation validation | 17-05-2022 | 02-03-2023 |
| W1309 | Female | Training, Millet & GPS validation | 13-05-2022 | 22-11-2022 |
| W1393 | Male | Training, Millet & GPS validation | 13-05-2022 | 02-05-2024 |
| W1413 | Male | Training, Millet & GPS validation | 25-02-2021 | 14-06-2021 |
| W1415 | Female | Training, Millet & GPS validation | 25-02-2021 | 30-01-2022 |
| W1746 | Male | Training, Millet & GPS validation | 11-02-2022 | 23-07-2024 |

| | | | | |
|---|---|---|---|---|
| W1869 | Male | Training, Millet & GPS validation | 05-11-2022 | 19-03-2023 |
| W2625 | Female | Training, Millet & GPS validation | 17-05-2022 | 23-08-2023 |
| W2875 | Male | Training, Millet & GPS validation | 09-02-2022 | 19-02-2023 |
| WT00043 | Male | Training, Millet & GPS validation | 17-05-2022 | 06-06-2023 |
| WT00162 | Male | Training, Millet & GPS validation | 24-03-2021 | 11-05-2021 |
| WT00500 | Female | Training, Millet & GPS validation | 25-02-2021 | 15-05-2024 |
| WT00584 | Male | Training, Millet & GPS validation | 17-05-2022 | 17-03-2023 |
| W1951 | Male | Night validation | 17-04-2023 | 24-09-2023 |
| W2885 | Female | Night validation | 25-05-2023 | 06-06-2023 |
| W2890 | Male | Night validation | 25-05-2023 | 06-06-2023 |
| W1286 | Male | Synchronisation validation | 02-06-2022 | 28-04-2024 |
| WT00584 | Male | Synchronisation validation | 17-05-2022 | 17-03-2023 |

Table S2: Ethogram used to label video data from n=14 birds.

| **Behaviour** | **Definition** |
|---|---|
| *Locomotion* (n=3) | |
| Walking | Slow speed travel |
| Trotting | Medium speed travel with ‘bounce’ |
| Running | Full speed travelling |
| *Foraging* (n=5) | |
| Pecking | Pecking downwards at the ground/short grass |
| Nipping | Pecking at head-level height at bush/tall grass/cactus |
| Searching | Slow speed walking with head down |

| | |
|---|---|
| Drinking | Drinking water by lowering head to water and then lifting head for water to run down oesophagus |
| Scratching | Scratching at the ground to dislodge earth |
| *Stationary behaviours* (n=4) | |
| Lying | Lying down on the side of the body, resting |
| Sitting | Sitting down, resting |
| Alert | Standing still, head up, looking around (vigilant). |
| Active sitting | Preening or pecking while in seated position |
| *Self-directed* (n=2) | |
| Preening | Preening feathers while standing up |
| Dust bathing | Sitting down in the dust, moving wings to coat body in dust |
| *Vocalisation* (n=1) | |
| *Long-distance call* | Loud call made while standing up, neck stretched out. |
| *Other* (n=2) | |
| Other | Behaviours that do not fall in the above categories (including: transitions between behaviours, agonistic interactions) |
| Not visible | When the bird is not clearly in view or obscured by object or another bird |

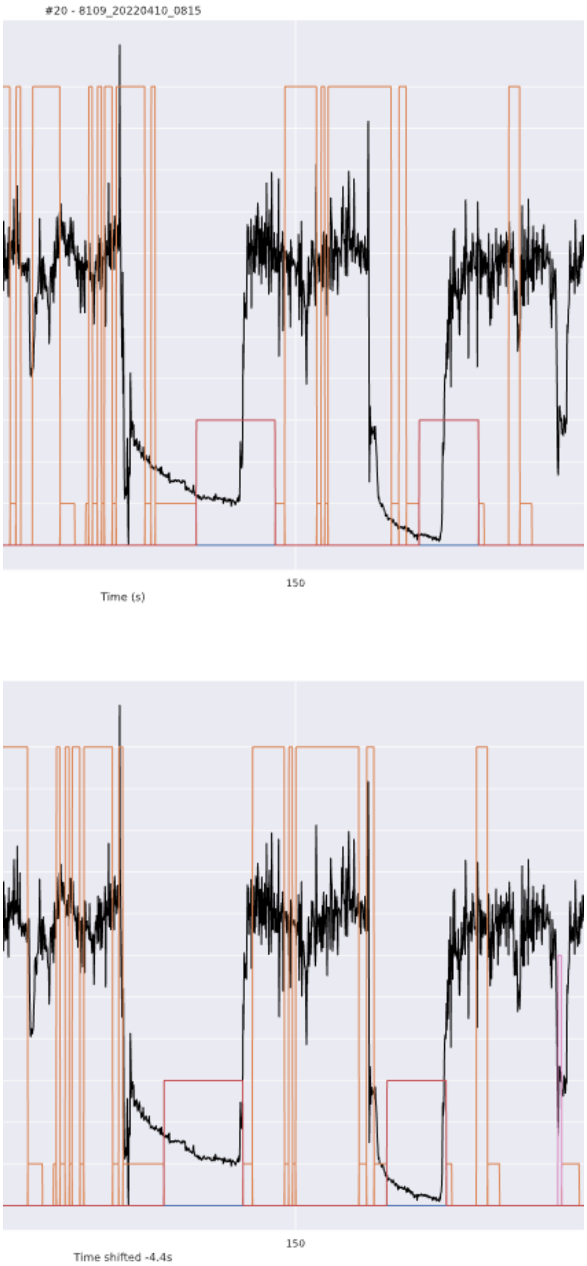


**Figure S1**: Example of time stamp correction when matching labelled video data to accelerometer signals. Here the black lines denote raw VeDBA with the orange tall boxes showing ‘feeding’ behaviour and red short boxes showing ‘alert’ behaviour. In the upper panel, the label is incorrectly matched to the acceleration signal. In the lower panel, the label is moved (- 4.4 seconds) to correctly align the acceleration signals with the behaviours.

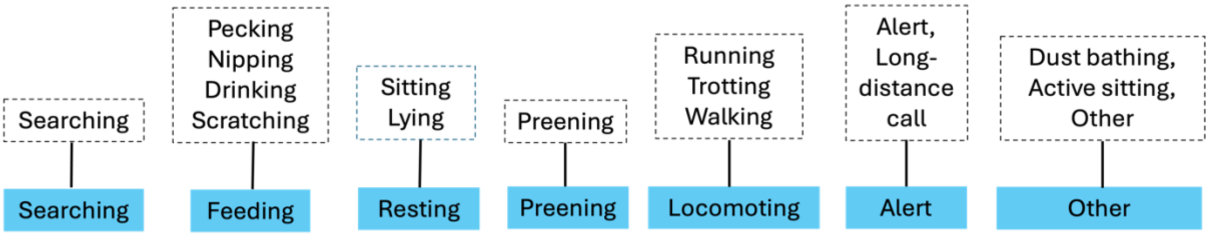


**Figure S2**: Merging of behavioural annotations from ethogram (top row) into behavioural categories which were fed into the random forest model (bottom row). Merging was based on similarity in biological function. Apart from 'Other' which represents the remaining categories that were not of biological interest (e.g., transitions) or had small sample sizes (e.g., dust bathing and social interactions).

**Table S3**: Behavioural labels and their associated sample size (in seconds), the number of independent events and the number of birds recorded performing this behaviour.

| Behaviour | Sample size (seconds) | # independent events | # individual birds |
|---|---|---|---|
| Alert | 5,086 | 542 | 14 |
| Feeding | 8,965 | 1,985 | 14 |
| Locomoting | 4,272 | 1,126 | 14 |
| Other | 7,973 | 2,775 | 14 |
| Preening | 6,098 | 512 | 14 |
| Searching | 6,369 | 2,035 | 14 |
| Sitting | 4,277 | 155 | 7 |

**Table S4: Confusion matrix of random forest model**: Predicted behaviours (rows) and observed behaviours (columns). Instances where the model correctly classified the behaviour (true positives) are in grey boxes, instances where the model incorrectly classified the behaviour (false positives) are in rows, instances where the model missed the correct behaviour (false negatives) are in columns.

| | Alert | Feeding | Locomoting | Other | Preening | Searching | Sitting |
|---|---|---|---|---|---|---|---|
| **Alert** | 68.3 | 0.1 | 2.9 | 15.0 | 5.1 | 0.1 | 8.5 |
| **Feeding** | 0.1 | 82.9 | 0.3 | 5.2 | 0.6 | 11.9 | 0.0 |
| **Locomoting** | 3.2 | 2.9 | 63.3 | 9.8 | 5.6 | 15.1 | 0.0 |
| **Other** | 9.7 | 4.7 | 6.6 | 48.1 | 14.4 | 9.3 | 7.1 |
| **Preening** | 3.1 | 1.7 | 4.9 | 17.9 | 68.6 | 2.7 | 1.1 |
| **Searching** | 0.3 | 12.5 | 8.0 | 11.4 | 1.5 | 66.4 | 0.1 |
| **Sitting** | 6.8 | 0.1 | 0.0 | 7.5 | 0.3 | 0.0 | 85.3 |

**Table S5: Confusion matrix of annotators**: Behaviours labelled by annotator 1 considered the 'true' behaviour (rows) and behaviours labelled by annotator(s) 2 (columns). Instances where the annotators are in agreement (true positives) are in grey boxes, instances where annotator(s) 2 incorrectly classified the behaviour (false positives) are in rows, instances where annotator(s) 2 missed the correct behaviour (false negatives) are in columns.

| | Alert | Feeding | Locomoting | Other | Preening | Searching | Sitting |
|---|---|---|---|---|---|---|---|
| **Alert** | 76.4 | 0.0 | 0.7 | 16.9 | 6.1 | 0.0 | 0.0 |
| **Feeding** | 0.2 | 97.0 | 0.2 | 2.2 | 0.0 | 0.5 | 0.0 |
| **Locomoting** | 0.0 | 0.7 | 81.7 | 8.2 | 0.2 | 9.3 | 0.0 |
| **Other** | 6.8 | 4.9 | 5.2 | 59.8 | 3.5 | 9.4 | 10.3 |
| **Preening** | 0.4 | 0.0 | 0.2 | 2.2 | 97.2 | 0.0 | 0.0 |
| **Searching** | 0.0 | 11.2 | 2.5 | 2.8 | 0.7 | 82.8 | 0.0 |
| **Sitting** | 0.0 | 0.0 | 0.0 | 2.7 | 0.0 | 0.0 | 97.3 |

**Table S6**: Results from simulation where the propensity of the behaviour of interest (alert, feeding, locomoting, preening, sitting, shown in the grey band within the table) is increased between two periods in the data by a given amount (simulated change). Detected mean change captures the mean of the change between the two periods over all 100 simulated studies. Proportion of P < 0.05 is the proportion of the simulated studies that detected a statistically significant change. Corresponding figure panels (see Fig. 2) are also given in the grey band.

| **Simulated change** | **Detected mean change** | **Proportion of *P* < 0.05** | **Detected mean change** | **Proportion of *P* < 0.05** |
|---|---|---|---|---|
| *Fig. 2b* | *Feeding (true behaviour)* | | *Searching (confused behaviour)* | |
| **0** | -0.001 | 0.2 | <0.001 | 0 |
| **0.0125** | 0.098 | 0.8 | -0.001 | 0 |
| **0.025** | 0.020 | 1.0 | <0.001 | 0.1 |
| **0.05** | 0.040 | 1.0 | -0.003 | 0.1 |
| **0.075** | 0.059 | 1.0 | -0.005 | 0.4 |
| **0.01** | 0.075 | 1.0 | -0.004 | 0.2 |
| **0.125** | 0.097 | 1.0 | -0.007 | 0.8 |
| **0.15** | 0.119 | 1.0 | -0.009 | 1.0 |
| **0.175** | 0.136 | 1.0 | -0.011 | 1.0 |
| **0.2** | 0.155 | 1.0 | -0.011 | 1.0 |
| *Fig. 2f* | *Sitting (true behaviour)* | | *Alert (confused behaviour)* | |
| **0** | <0.001 | 0.01 | <0.001 | 0 |
| **0.0125** | 0.010 | 1.0 | <0.001 | 0 |
| **0.025** | 0.021 | 1.0 | <0.001 | 0.1 |
| **0.05** | 0.042 | 1.0 | -0.002 | 0.2 |
| **0.075** | 0.061 | 1.0 | -0.003 | 0.1 |
| **0.01** | 0.083 | 1.0 | -0.004 | 0.3 |
| **0.125** | 0.103 | 1.0 | -0.007 | 0.6 |
| **0.15** | 0.124 | 1.0 | -0.009 | 0.9 |
| **0.175** | 0.146 | 1.0 | -0.008 | 1.0 |

| | | | | |
|---|---|---|---|---|
| **0.2** | 0.165 | 1.0 | -0.009 | 1.0 |
| *Fig. 2a* | *Alert (true behaviour)* | | *Other (confused behaviour)* | |
| **0** | <0.001 | 0.1 | <0.001 | 0 |
| **0.0125** | 0.008 | 0.7 | <0.001 | 0 |
| **0.025** | 0.017 | 1.0 | -0.002 | 0.1 |
| **0.05** | 0.032 | 1.0 | -0.001 | 0.1 |
| **0.075** | 0.049 | 1.0 | -0.003 | 0.3 |
| **0.01** | 0.065 | 1.0 | -0.004 | 0.3 |
| **0.125** | 0.081 | 1.0 | -0.006 | 0.3 |
| **0.15** | 0.096 | 1.0 | -0.006 | 0.4 |
| **0.175** | 0.114 | 1.0 | -0.008 | 0.8 |
| **0.2** | 0.130 | 1.0 | -0.011 | 0.8 |
| *Fig. 2c* | *Locomoting (true behaviour)* | | *Searching (confused behaviour)* | |
| **0** | <0.001 | 0 | <0.001 | 0.1 |
| **0.0125** | 0.009 | 1 | 0.001 | 0.2 |
| **0.025** | 0.015 | 1 | -0.001 | 0 |
| **0.05** | 0.030 | 1 | -0.001 | 0 |
| **0.075** | 0.044 | 1 | <0.001 | 0 |
| **0.01** | 0.056 | 1 | -0.002 | 0.1 |
| **0.125** | 0.073 | 1 | -0.002 | 0.1 |
| **0.15** | 0.090 | 1 | -0.002 | 0.1 |
| **0.175** | 0.104 | 1 | <0.001 | 0 |
| **0.2** | 0.118 | 1 | <0.001 | 0.1 |
| *Fig. 2d* | *Preening (true behaviour)* | | *Other (confused behaviour)* | |
| **0** | <0.001 | 0.2 | <0.001 | 0 |
| **0.0125** | 0.009 | 0.9 | -0.001 | 0.1 |
| **0.025** | 0.014 | 1.0 | <0.001 | 0.2 |
| **0.05** | 0.032 | 1.0 | -0.001 | 0 |
| **0.075** | 0.047 | 1.0 | 0.002 | 0 |
| **0.01** | 0.064 | 1.0 | -0.003 | 0.2 |

| | | | | |
|---|---|---|---|---|
| **0.125** | 0.080 | 1.0 | <0.001 | 0.1 |
| **0.15** | 0.096 | 1.0 | -0.001 | 0 |
| **0.175** | 0.112 | 1.0 | -0.003 | 0.2 |
| **0.2** | 0.128 | 1.0 | -0.005 | 0.3 |

**A worked example of the effect of mislabels on training and evaluating models.**

Errors in labelling training data can have different impacts on the real and estimated performance of a model used for a classification task. Here we introduce three types of errors into empirical data comprising classification of three species of birds: nuthatches, great tits, blue tits. Pictures were collected at artificial feeding stations and pre-processed (e.g. removing the background to select only the pixels corresponding to the birds) using the methodology described in Ferreira *et al.* (2020b). The three types of error consist of different types of erroneous labels, which are used for both training and validation dataset (see supplementary material for Box 1):

*Uniform error*: All pictures from all three species of birds have an equal chance of being mislabelled. This illustrates a situation in which all classes (i.e. species of birds in this example) are equally difficult to label.

*Class-dependent error*: Pictures from nuthatches are always correctly labelled, but pictures of great tits could be mislabelled as blue tits and *vice-versa*. This error type represents a situation where some classes are easy to classify, but others are more difficult to distinguish.

*Class- and feature-dependent error*: Here we artificially introduce a feature to the pictures and correlate this feature with the probability of mislabel. For all pictures of great tits and blue tits that had a wrong label we also blurred those images. This would simulate a situation in which the human labeller would have great difficulties in classifying these two species of birds but only when the image quality was low (in this case blur).

For all types of error, we train several models (a VGG19 backbone model pre-trained on ImageNet dataset, see Supplementary material for details) that vary in the proportion of mislabels (equal for training and validation dataset) and evaluate each model using simple accuracy (N instances of predicted labels matching the annotated label/ all instances) on the validation dataset (reported accuracy). We then compare the performance to the same set of validation images but without any error in the labels, representing the true performance of the models (real accuracy).

For all cases we can see that the reported accuracy is misleading when error is present in the labels. For both uniform (a) and class-dependent error (b), the model is able to learn beyond human capacity (i.e. makes fewer errors in predicting labels than the proportion of mislabels in the training data). In this scenario, which is likely to be common in ecological research, standard metrics would underestimate the model's true performance—and this underestimation increases linearly with the extent of the error. For the class- and feature-base error (c), we would overestimate the model's ability to generalize to unseen data. This is likely because the model learns to associate the interaction between blur and pictures of great tits or blue tits, classifying all blurred pictures with the label of the other species. Since this error is also present in the validation dataset the model's accuracy is always reported as being high while the true generalization capability of the model keeps degrading. This result suggests that it may be better practice to avoid including data that are ambiguous in the validation dataset. Finally, it is important to note that real-world error is often complex and all types of errors as well as other variations to the ones mentioned here can be present in a single dataset.

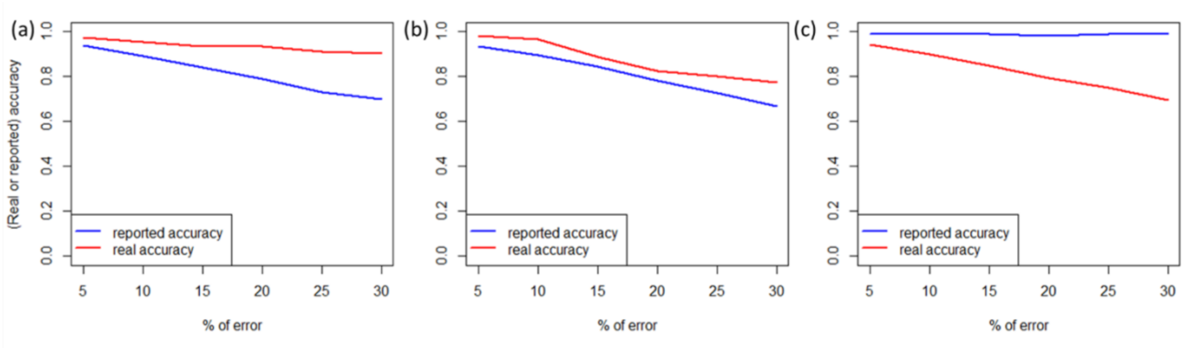


### Training details

For the models we used 5 different individual birds per species (15 in total), 2500 pics for training per species (7500 total) 500 for validation per species (1500 total). All images were resized to 224x224 pixels. For all types of errors, we used the same model structure. A VGG19 architecture pre-trained on ImageNet dataset for which the top classification layer was replaced by a dense layer with 256 neurons followed by a

Softmax classification layer with 3 neuros (each representing one of the three species). Models were trained with Adam optimizer, a learning rate of 1e-5 and a batch size of 30. All models were trained until there was no further improved on the validation loss for more than five consecutive epochs. All models were trained using python implementation of TensorFlow. See training scripts for more details.

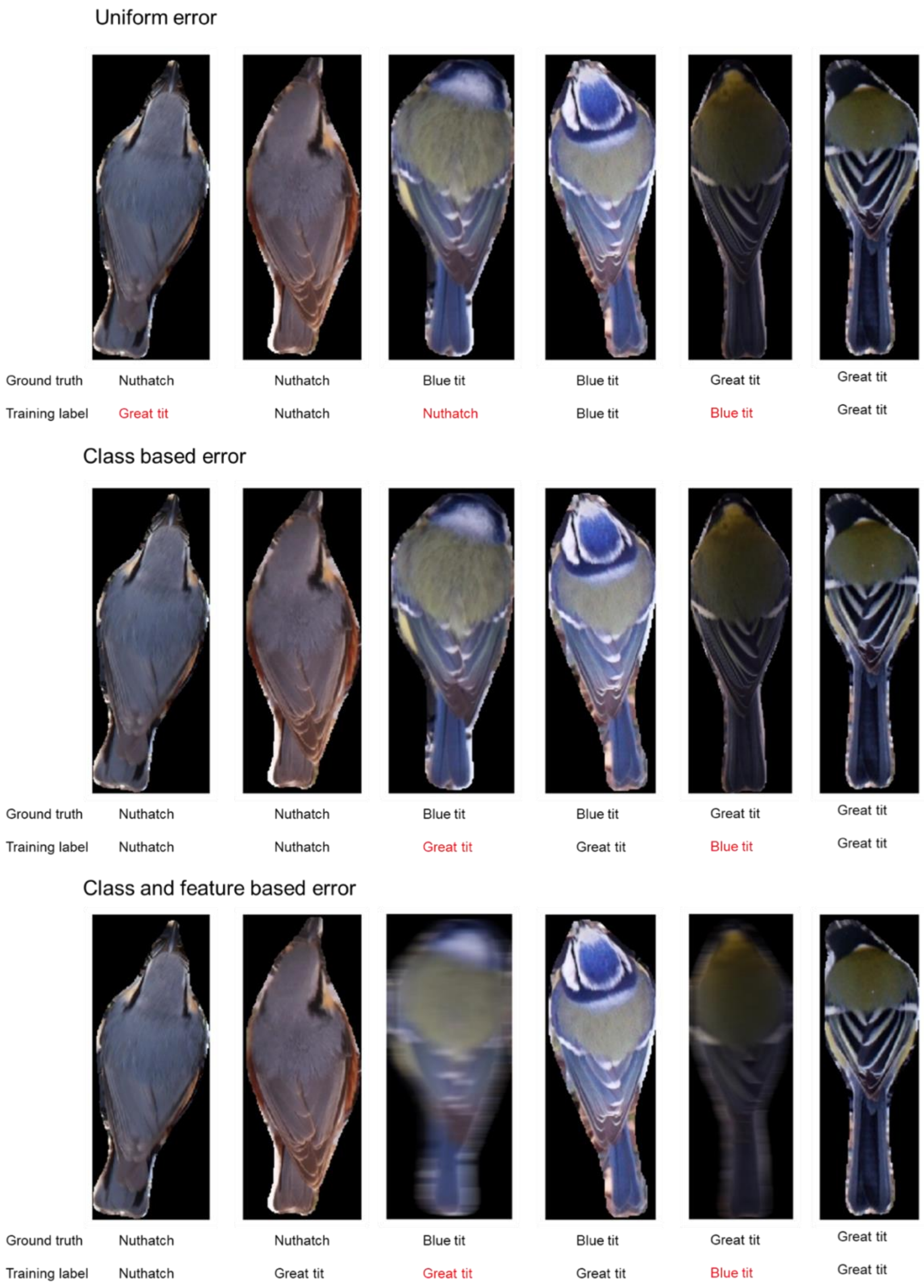


**Figure S3**– Illustrative examples of each type of simulated error used in ‘A worked example of the effect of mislabels on training and evaluating models’: uniform error (first row), class based error (middle row) and class and feature based error (bottom row).

127
